# A prefusion SARS-CoV-2 spike RNA vaccine is highly immunogenic and prevents lung infection in non-human primates

**DOI:** 10.1101/2020.09.08.280818

**Authors:** Annette B. Vogel, Isis Kanevsky, Ye Che, Kena A. Swanson, Alexander Muik, Mathias Vormehr, Lena M. Kranz, Kerstin C. Walzer, Stephanie Hein, Alptekin Güler, Jakob Loschko, Mohan S. Maddur, Kristin Tompkins, Journey Cole, Bonny G. Lui, Thomas Ziegenhals, Arianne Plaschke, David Eisel, Sarah C. Dany, Stephanie Fesser, Stephanie Erbar, Ferdia Bates, Diana Schneider, Bernadette Jesionek, Bianca Sänger, Ann-Kathrin Wallisch, Yvonne Feuchter, Hanna Junginger, Stefanie A. Krumm, André P. Heinen, Petra Adams-Quack, Julia Schlereth, Christoph Kröner, Shannan Hall-Ursone, Kathleen Brasky, Matthew C. Griffor, Seungil Han, Joshua A. Lees, Ellene H. Mashalidis, Parag V. Sahasrabudhe, Charles Y. Tan, Danka Pavliakova, Guy Singh, Camila Fontes-Garfias, Michael Pride, Ingrid L. Scully, Tara Ciolino, Jennifer Obregon, Michal Gazi, Ricardo Carrion, Kendra J. Alfson, Warren V. Kalina, Deepak Kaushal, Pei-Yong Shi, Thorsten Klamp, Corinna Rosenbaum, Andreas N. Kuhn, Özlem Türeci, Philip R. Dormitzer, Kathrin U. Jansen, Ugur Sahin

## Abstract

To contain the coronavirus disease 2019 (COVID-19) pandemic, a safe and effective vaccine against the new severe acute respiratory syndrome coronavirus-2 (SARS-CoV-2) is urgently needed in quantities sufficient to immunise large populations. In this study, we report the design, preclinical development, immunogenicity and anti-viral protective effect in rhesus macaques of the BNT162b2 vaccine candidate. BNT162b2 contains an LNP-formulated nucleoside-modified mRNA that encodes the spike glycoprotein captured in its prefusion conformation. After expression of the BNT162b2 coding sequence in cells, approximately 20% of the spike molecules are in the one-RBD ‘up’, two-RBD ‘down’ state. Immunisation of mice with a single dose of BNT162b2 induced dose level-dependent increases in pseudovirus neutralisation titers. Prime-boost vaccination of rhesus macaques elicited authentic SARS-CoV-2 neutralising geometric mean titers 10.2 to 18.0 times that of a SARS-CoV-2 convalescent human serum panel. BNT162b2 generated strong T_H_1 type CD4^+^ and IFNγ^+^ CD8^+^ T-cell responses in mice and rhesus macaques. The BNT162b2 vaccine candidate fully protected the lungs of immunised rhesus macaques from infectious SARS-CoV-2 challenge. BNT162b2 is currently being evaluated in a global, pivotal Phase 2/3 trial (NCT04368728).

## Introduction

Due to the shattering impact of the coronavirus disease 2019 (COVID-19) pandemic on human health and society, multiple collaborative research programs have been launched, leading to new insights and progress towards vaccine development. Soon after it emerged in December 2019, severe acute respiratory syndrome coronavirus-2 (SARS-CoV-2) was identified as a β-coronavirus with high sequence similarity to bat-derived SARS-like coronaviruses^1,2^. The globalised response is mirrored by the upload of over 92,000 viral genome sequences as of August 29, 2020, to GISAID (Global Initiative on Sharing All Influenza Data).

The trimeric spike glycoprotein (S) of SARS-CoV-2 binds its cellular receptor, angiotensin converting enzyme 2 (ACE2), through a receptor-binding domain (RBD), which is part of its N-terminal furin cleavage fragment (S1)^3,4^. S rearranges to translocate the virus into cells by membrane fusion^5,6^. The C-terminal furin cleavage fragment (S2) contains the fusion machinery^7^. Membrane fusion can be blocked by mutating S residues 986 and 987 to prolines, producing an S antigen stabilised in the prefusion conformation (P2 S)^8–10^. The RBD is a key target for virus neutralising antibodies, with an ‘up’ conformation, in which more neutralising epitopes are exposed, and a ‘down’ conformation in which many epitopes are buried^5,10–12^. In addition, some neutralising antibodies bind S epitopes outside the RBD.

During this pandemic, fast vaccine availability is critical. COVID-19 vaccine candidates based on different platforms are already in clinical trials, with the most advanced based on viral vector and nucleic acid technologies^13–16^. We report the preclinical development of BNT162b2, a lipid-nanoparticle (LNP) formulated N^1^-methyl-pseudouridine (m1Ψ) nucleoside-modified mRNA (modRNA) vaccine candidate that encodes P2 S with a native furin cleavage site resulting in the S1 and S2 cleavage fragments (Fig. 1a). The m1Ψ-modification dampens innate immune sensing, and, together with optimised non-coding sequence elements, increases RNA translation *in vivo*^17–19^. ModRNA vaccines have already proven immunogenic for several viral targets^20,21^. BNT162b2 has been evaluated in phase 1 clinical trials in the US (NCT04368728) and Germany (NCT04380701, EudraCT: 2020-001038-36), and is now being evaluated in a pivotal, global, phase 2/3 safety and efficacy study^15^.

**Figure 1.**
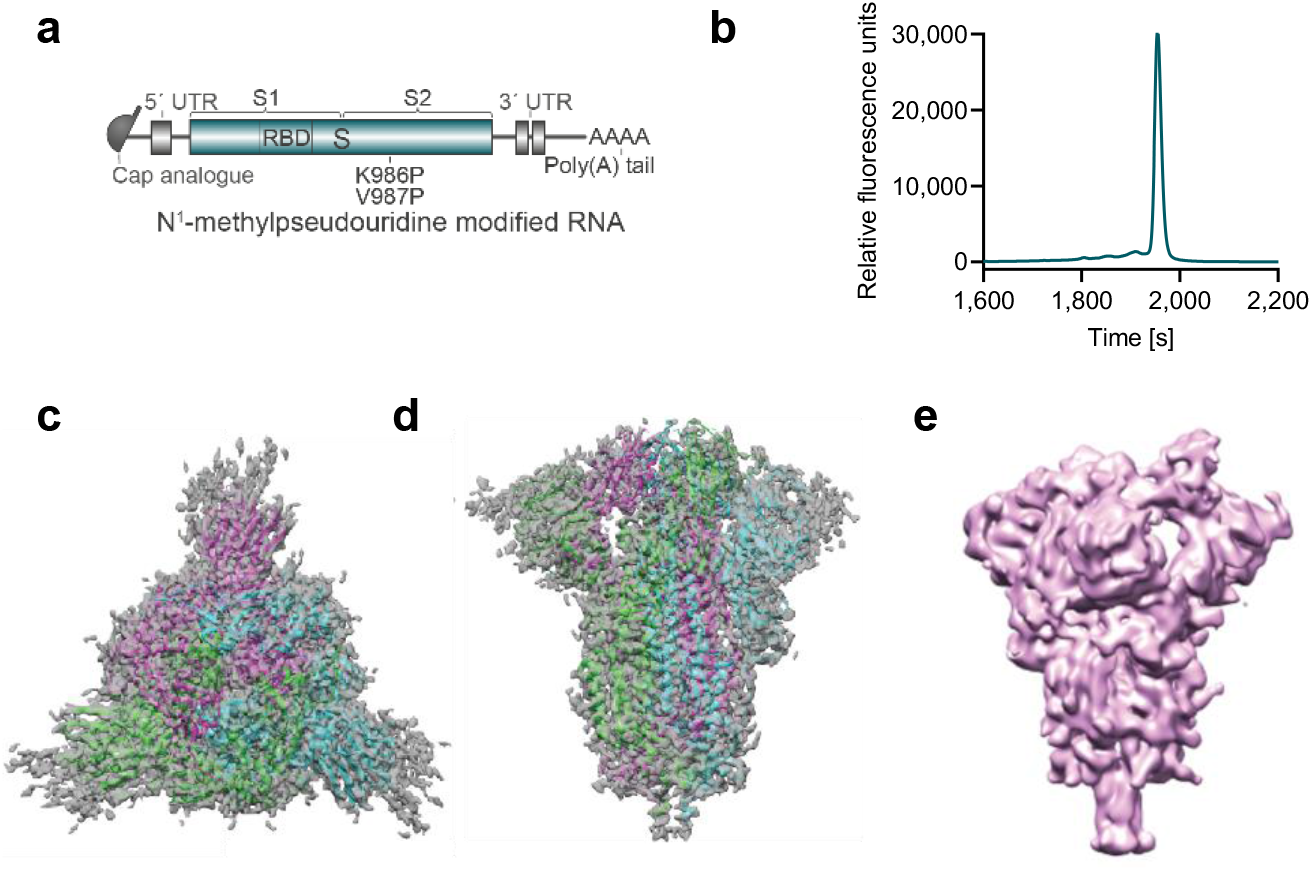
Vaccine design and characterisation of the expressed antigen. **a**, BNT162b2 RNA structure. UTR, untranslated region; S, SARS-CoV-2 S glycoprotein; S1, N-terminal furin cleavage fragment; S2, C-terminal furin cleavage fragment; RBD, receptor-binding domain. Positions of the P2 mutation (K986P and V897P) are indicated. **b**, Liquid capillary electropherogram of *in vitro* transcribed BNT162b2 RNA. **c**, A 3.29 Å cryoEM map of P2 S, with fitted and refined atomic model, viewed down the three-fold axis toward the membrane. **d,** Cryo-EM map and model of (d) viewed perpendicular to the three-fold axis. **e,** Mass density map of TwinStrep-tagged P2 S produced by 3D classification of images extracted from cryo-EM micrographs with no symmetry averaging. This class, in the one-RBD ‘up’, two RBD ‘down’ positioning, represents 20.4% of the population.

## Results

BNT162b2 RNA *in vitro* transcribed by T7 polymerase from a plasmid DNA template has a single, sharp-peak microfluidic capillary electrophoresis profile, consistent with its calculated length of 4,283 nucleotides, indicating purity and integrity (Fig. 1b). When HEK293T/17 cells were incubated with BNT162b2 (which is LNP-formulated) or with BNT162b2 RNA mixed with a transfection reagent, robust expression of P2 S was detectable by flow cytometry (Extended Data Fig. 1a).

For structural characterisation, P2 S was expressed in Expi293F cells from DNA that encodes the same amino acid sequence as BNT162b2 RNA, with the addition of a C-terminal TwinStrep tag for affinity purification. The trimeric P2 S bound the human ACE2 peptidase domain (PD), and an anti-RBD human neutralising antibody B38 with high affinity (K_D_ 1 nM, Extended Data Fig. 1b,c)^22^. Structural analysis by cryo-electron microscopy (cyro-EM) produced a 3.29 Å nominal resolution mass density map, into which a previously published atomic model^10^ was fitted and rebuilt (Fig. 1c,d; Extended Data Fig. 2, Extended Data Table 1). The rebuilt model shows good agreement with reported structures of prefusion full-length wild type S and its ectodomain with P2 mutations^5,10^. Three-dimensional classification of the dataset showed a class of particles that was in the one RBD ‘up’ (accessible for receptor binding), two RBD ‘down’ (closed) conformation and represented 20.4% of the trimeric molecules. The remainder were in the all RBD ‘down’ conformation (Fig. 1e, Extended Data Fig. 2c). The RBD in the ‘up’ conformation was less well resolved than other parts of the structure, suggesting conformational flexibility and a dynamic equilibrium between RBD ‘up’ and RBD ‘down’ states as also suggested by others^5,23^. Nevertheless, the binding and structural analyses indicate that the BNT162b2 RNA sequence encodes a recombinant P2 S that authentically presents the ACE2 binding site and other epitopes targeted by SARS-CoV-2 neutralising antibodies.

To characterise BNT162b2-elicited B- and T-cell responses, BALB/c mice were immunized intramuscularly (IM) once with 0.2, 1, or 5 μg BNT162b2 or received a buffer control. S1- and RBD-binding serum IgG developed rapidly at all dose levels in a dose-dependent manner. For S1-binding antibodies, the geometric mean concentration (GMC) in the 5 μg group was 386 μg/mL at Day 28 (Fig. 2a, Extended Data Fig. 3a). At Day 28 after immunisation, vaccine-elicited IgG had a strong binding affinity for S1 (geometric mean K_D_ 12 nM) and the RBD (geometric mean K_D_ 0.99 nM), with both having a low off-rate (Extended Data Fig. 3b). SARS-CoV-2 neutralising activity in mouse serum was measured by a vesicular stomatitis virus (VSV)-based SARS-CoV-2 pseudovirus neutralisation assay. Fifty percent pseudovirus neutralisation geometric mean titers (pVNT_50_ GMTs) increased steadily after immunisation to 26, 176, and 296 on Day 28 for the 0.2, 1, and 5 μg dose levels, respectively (Fig. 2b, Extended Data Fig. 3c).

**Figure 2.**
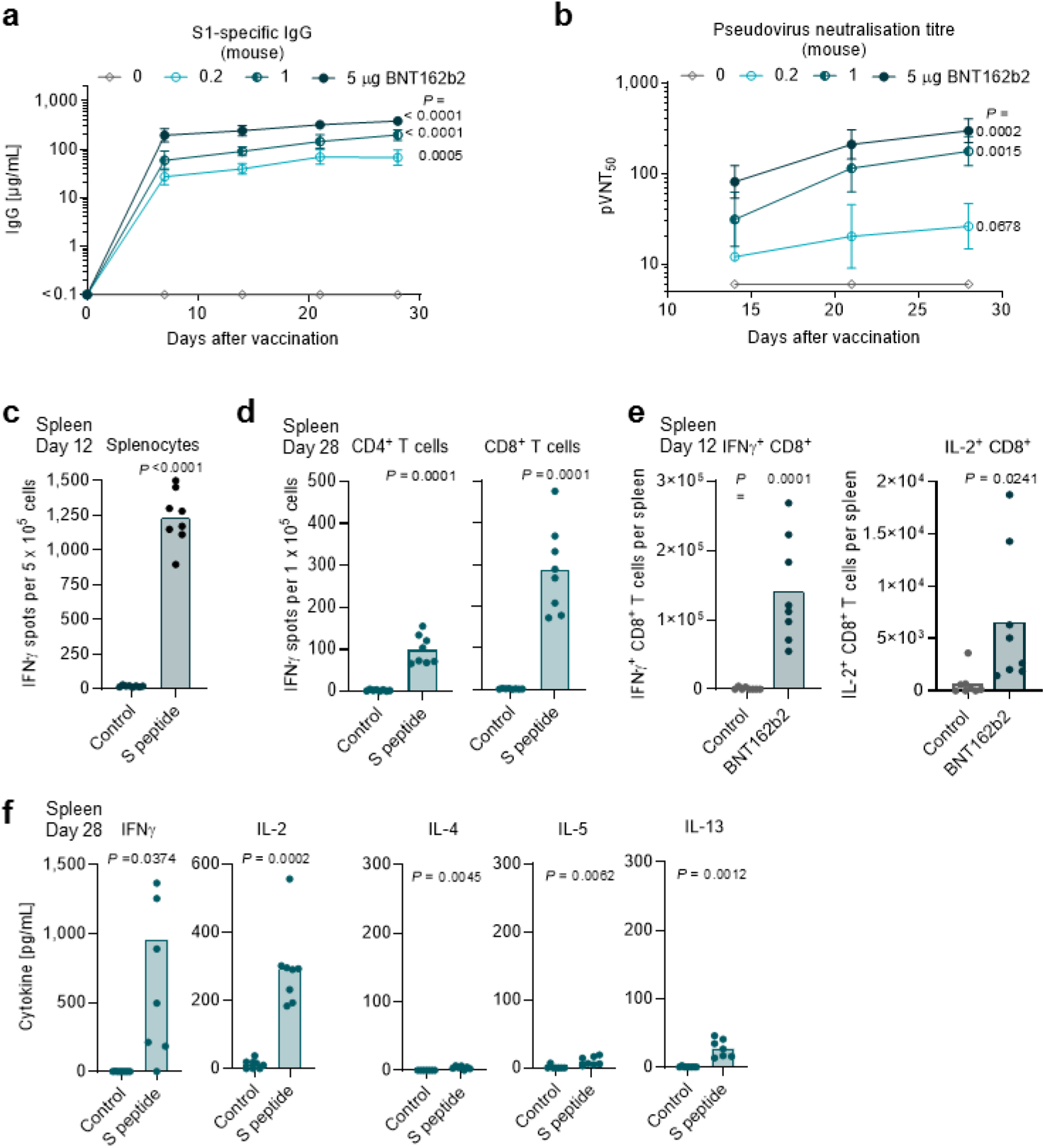
Mouse immunogenicity. BALB/c mice (n=8 per group unless otherwise specified) were immunised intramuscularly (IM) with a single dose of with 0.2, 1 or 5 μg BNT162b2 or buffer. Geometric mean of each group ± 95% CI, P-values compare Day 28 to non-immunised (0 μg; n=8) baseline sera (multiple comparison of mixed-effect analysis using Dunnett’s multiple comparisons test) (a, b). **a**, S1-binding IgG responses in sera obtained 7, 14, 21 and 28 days after immunisation with 0, 0.2, 1, or 5 μg BNT162b2, determined by ELISA. For day 0 values, a pre-screening of randomly selected animals was performed (n=4). **b,** VSV-SARS-CoV-2 pseudovirus 50% serum neutralising titers (pVNT_50_) of sera from (a). **c-f,** Splenocytes of BALB/c mice immunised IM with BNT162b2 or buffer (control) were *ex vivo* re-stimulated with full-length S peptide mix or negative controls ([c], [e], [f]: no peptide; [d]: irrelevant peptide). Individual values and mean of each group, P-values were determined by a two-tailed paired t-test. **c**, IFNγ ELISpot of splenocytes collected 12 days after immunisation with 5 μg BNT162b2. **d**, IFNγ ELISpot of isolated splenic CD4^+^ or CD8^+^ T cells collected 28 days after immunisation with 1 μg BNT162b2. **e**, CD8^+^ T-cell specific cytokine release by splenocytes collected 12 days after immunisation with 5 μg BNT162b2 or buffer (Control), determined by flow cytometry. S-peptide specific responses are corrected for background (no peptide). **f**, Cytokine production by splenocytes collected 28 days after immunisation with 1 μg BNT162b2, determined by bead-based multiplex analysis (n=7 for IL-4, IL-5 and IL-13, one outlier removed via routs test [Q=1%] for the S peptide stimulated samples).

**Figure 3.**
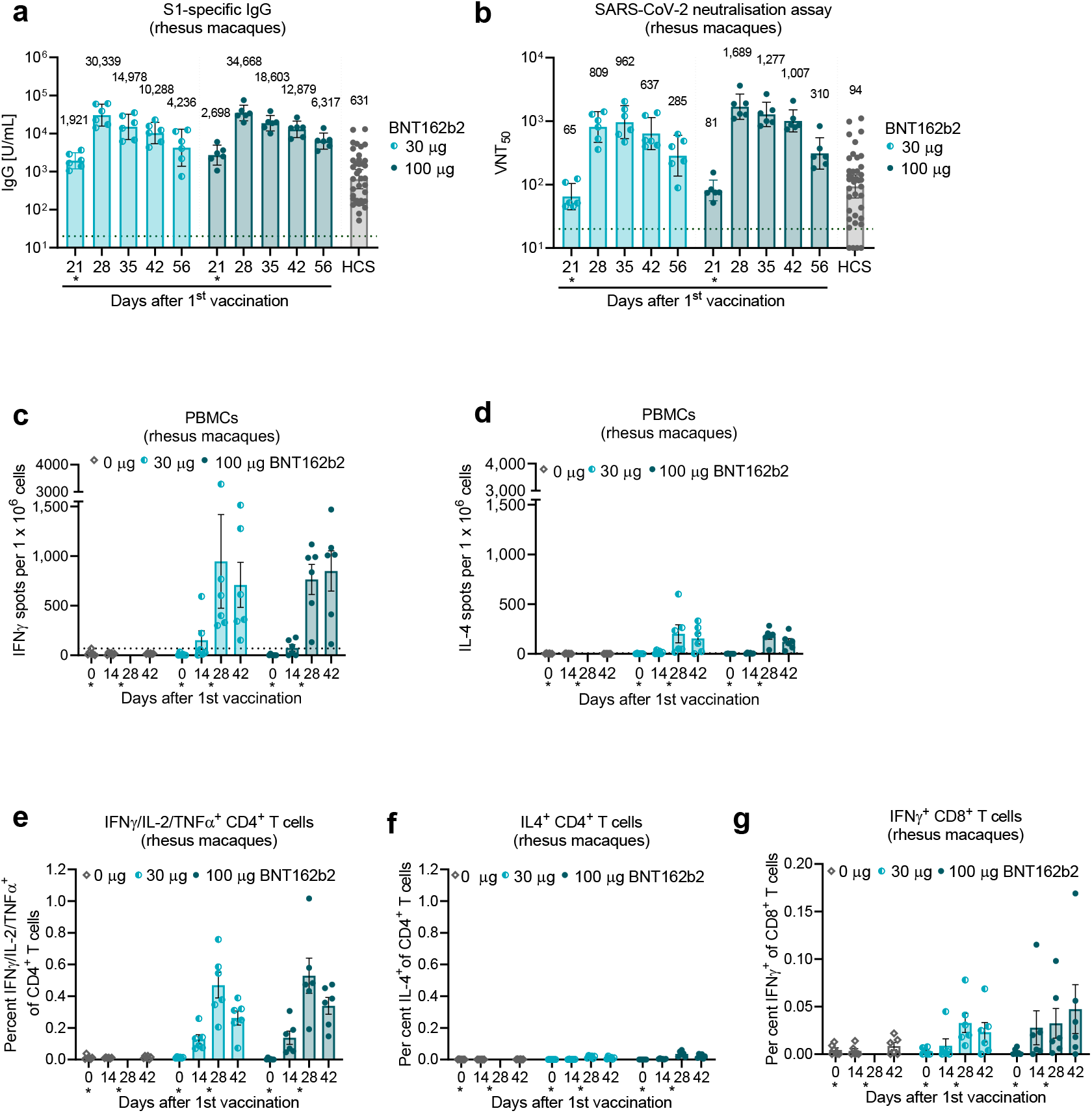
Rhesus macaque immunogenicity. Rhesus macaques (n=6 per group) were immunised on Days 0 and 21 with 30 μg or 100 μg BNT162b2 or buffer. Sera and PBMCs were collected at the times indicated. Human convalescent sera (HCS) were obtained from SARS-CoV-2-infected patients at least 14 days after PCR-confirmed diagnosis and at a time when acute COVID-19 symptoms had resolved (n=38). **a**, Concentration, in arbitrary units, of IgG binding recombinant SARS-CoV-2 S1. **b**, SARS-CoV-2 50% virus neutralisation titers (VNT_50_). **c-g**, PBMCs collected on Day 42 were *ex vivo* re-stimulated with full-length S peptide mix. **c,** IFNγ, and **d,** IL-4 ELISpot. **e**, **f**, CD4^+^ T-cell specific, and **g**, CD8^+^ T-cell specific cytokine release, determined by flow cytometry. Heights of bars indicate the geometric (a-b) or arithmetic (c-g) means for each group. Whiskers indicate 95% confidence intervals (CI’s; a-b) or standard errors of means (SEMs; c-g). Every symbol represents one animal. Horizontal dotted lines mark the LLODs. Values below the LLOD set to ½ the LLOD. Asterisks below the x-axis indicate the day of Dose 2.

A high fraction of splenocytes of CD4^+^ and CD8^+^ T-cell phenotype isolated from mice on Days 12 and 28 after BNT162b2-immunisation had a strong antigen-specific IFNγ and IL-2 response in ELISpot and intracellular cytokine staining flow cytometry analysis when re-stimulated *ex vivo* with a full-length S peptide pool (Fig. 2c-e). Total splenocytes harvested on Day 28 and *ex vivo* re-stimulated with the full-length S peptide pool secreted high levels of the T_H_1 cytokines IL-2 or IFNγ, but minute amounts of the T_H_2 cytokines IL-4, IL-5 and IL-13 as measured in multiplex immunoassays (Fig. 2f).

BNT162b2-induced effects on proliferation and dynamics of immune cell populations were assessed in injection site draining lymph nodes (dLNs), which are the principal immune-educated compartments for proficient T- and B-cell priming, and in blood and spleen for evaluation of its systemic effects. Higher numbers of plasma cells, class switched IgG1- and IgG2a-positive B cells, and germinal center B cells were observed in dLNs and spleens of mice 12 days after immunisation with 5 μg BNT162b2 than after immunisation with buffer (Extended Data Fig. 4a, b). In Day 7 post-immunisation blood, there were significantly fewer circulating B cells than in blood from buffer-immunised mice (Extended Data Fig. 4c), which may imply that B-cell homing to lymphoid compartments augments B cell counts in dLN and spleen. The dLNs from BNT162b2-immunised mice also have significantly elevated counts of CD8^+^ and CD4^+^ T cells, which was most pronounced for T follicular helper (T_FH_) cells, including ICOS^+^ subsets essential for germinal center formation (Extended Data Fig. 4a)^24^. BNT162b2 immunisation increased CD8^+^ T cell counts in the blood and T_FH_ cell counts in the spleen and blood (Extended Data Fig. 4b, c). These data indicate that BNT162b2 concurrently elicits strong SARS-CoV-2 pseudovirus neutralising titers and systemic T_H_1-driven CD4^+^ and CD8^+^ T-cell responses.

To assess BNT162b2-mediated protection in non-human primates, groups of six male, 2-4 year old rhesus macaques were immunised IM with 30 or 100 μg of BNT162b2 or saline control on Days 0 and 21. S1-binding IgG was readily detectable by Day 21 after Dose 1, and levels increased further after Dose 2 through Day 28 (Fig. 3a). Seven days after Dose 2 (Day 28), the GMCs of S1-binding IgG were 30,339 units (U)/mL (30 μg dose level) and 34,668 U/mL (100 μg dose level). For comparison, the S1-binding IgG GMC of a panel of 38 SARS-CoV-2 convalescent human sera was 631 U/mL, substantially lower than the GMCs of the immunised rhesus macaques after one or two doses.

Fifty percent virus neutralisation GMTs, measured by an authentic SARS-CoV-2 neutralisation assay^25^, were detectable in rhesus macaque sera by Day 21 after Dose 1 and peaked at a GMT of 962 (Day 35, 14 days after Dose 2 of 30 μg) or 1,689 (Day 28, 7 days after Dose 2 of 100 μg; Fig. 3b). Robust GMTs of 285 for 30 μg and 310 for 100 μg dose levels persisted to at least Day 56 (most recent time point tested). For comparison, the neutralisation GMT of the human convalescent serum panel was 94.

S-specific T-cell responses were analysed in BNT162b2-immunised rhesus macaques and saline-immunised controls by ELISpot and intracellular cytokine staining (ICS). Peripheral blood mononuclear cells (PBMCs) were collected before immunisation and at the times indicated after Doses 1 and 2. Strong IFNγ but minimal IL-4 responses were detected by ELISpot after Dose 2 (Fig. 3c,d, Extended Data Fig. 5). ICS confirmed that BNT162b2 elicited strong S-specific IFNγ producing T-cell responses, including a high frequency of CD4^+^ T cells that produced IFNγ, IL-2, and TNF but a low frequency of CD4^+^ T cells that produced IL-4, indicating a T_H_1-biased response (Fig. 3e,f). BNT162b2 also elicited S-specific IFNγ^+^ producing CD8^+^ T cells (Fig. 3g).

Six rhesus macaques that had received two immunisations with 100 μg BNT162b2 and three age-matched macaques that had received saline were challenged 55 days after Dose 2 with 1.05 × 10^6^ plaque forming units of SARS-CoV-2 (strain USA-WA1/2020), split equally between intranasal and intratracheal routes, as previously described^26^. Three additional non-immunised, age-matched rhesus macaques (sentinels) were mock-challenged with cell culture medium. Nasal and oropharyngeal (OP) swabs were collected and bronchoalveolar lavage (BAL) was performed at the times indicated, and samples were tested for SARS-CoV-2 RNA (genomic RNA or subgenomic transcripts) by reverse-transcription quantitative polymerase chain reaction (RT-qPCR; Fig. 4). All personnel performing clinical, radiological, histopathological, or RT-qPCR evaluations were blinded to the group assignments of the macaques.

**Figure 4.**
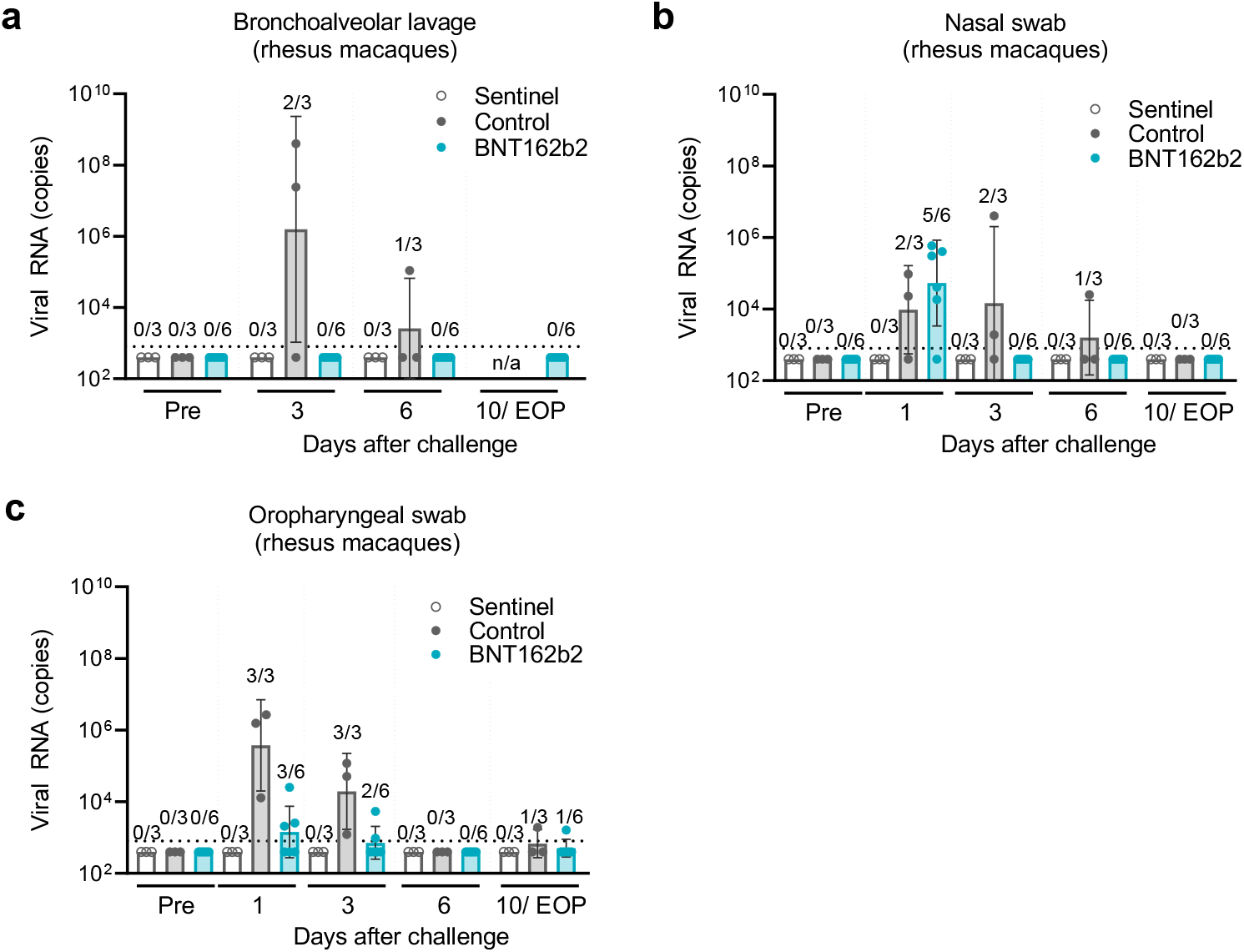
Protection of rhesus macaques from challenge with infectious SARS-CoV-2. Fifty-five days after the Dose 2 of 100 μg BNT162b2 (n=6) or saline control (n=3), rhesus macaques were challenged with 1.05 × 10^6^ total pfu of SARS-CoV-2 split equally between the IN and IT routes. Non-immunised rhesus macaques (n=3) were mock-challenged with cell culture medium (sentinel). Viral RNA levels were detected by RT-qPCR. BAL was performed and nasal and oropharyngeal (OP) swabs obtained at the indicated time points. Final collection of samples was on Day 10 relative to challenge for the sentinel and control groups and at the end of protocol (EOP) on Day 7 or 8 for the BNT162b2-immunised group. Ratios above data points indicate the number of viral RNA positive animals among all animals per group. Heights of bars indicate geometric means. Whiskers indicate geometric standard deviations. Every symbol represents one animal. Dotted lines indicate the lower limits of detection (LLOD). Values below the LLOD were set to ½ the LLOD. **a**, Viral RNA in bronchoalveolar lavage (BAL) fluid. **b**, Viral RNA in nasal swabs. **c**, Viral RNA in OP swabs. The statistical significance by a non-parametric test (Friedman’s test) of differences in viral RNA detection between control-immunised and BNT162b2-immunised animals after challenge was p=0.0014 for BAL fluid, p=0.2622 for nasal swabs, and p=0.0007 for OP swabs. n/a – not available.

Viral RNA was detected in BAL fluid from 2 of the 3 control-immunised macaques on Day 3 after challenge and from 1 of 3 on Day 6 (Fig. 4a). At no time point sampled was viral RNA detected in BAL fluid from the BNT162b2-immunised and SARS-CoV-2 challenged macaques (Fig. 4a). The difference in viral RNA detection in BAL fluid between BNT162b2-immunised and control-immunised rhesus macaques after challenge is highly statistically significant (by a nonparametric test, p=0.0014).

From control-immunised macaques, viral RNA was detected in nasal swabs obtained on Days 1, 3, and 6 after SARS-CoV-2 challenge; from BNT162b2-immunised macaques, viral RNA was detected only in nasal swabs obtained on Day 1 after challenge and not in swabs obtained on Day 3 or subsequently (Fig. 4b). The pattern of viral RNA detection from OP swabs was similar to that for nasal swabs (Fig. 4c).

In general, virus-challenged animals showed no clinical signs of significant disease. We conclude that the 2-4 year old male rhesus macaque challenge model is primarily a SARS-CoV-2 infection model and not a COVID-19 disease model.

## Discussion

We demonstrate that BNT162b2, an LNP-formulated, m1Ψ nucleoside-modified mRNA encoding SARS-CoV-2 S captured in a prefusion conformation is highly immunogenic in mice and rhesus macaques. Expression from DNA of protein with the BNT162b2-encoded amino acid sequence confirmed the P2 S prefusion conformation by cryo-EM. This analysis confirmed that the antigenically important RBD can assume the ‘up’ conformation, with the receptor binding site, rich in neutralising epitopes, accessible in a proportion of the molecules^27^. The alternative states observed likely reflect a dynamic equilibrium between RBD ‘up’ and ‘down’ positions^10,23^. Binding of expressed and purified P2 S to ACE2 and a neutralising monoclonal antibody further demonstrates its conformational and antigenic integrity.

In mice, a single injection of BNT162b2 elicited high neutralizing titers and strong T_H_1 and T_FH_ type CD4^+^ and IFNγ^+^IL-2^+^ CD8^+^ T-cell responses. Both BNT162b2 induced CD4^+^ T-cell types may support antigen-specific antibody generation and maturation, and potentially protection from infectious challenge. Limitation and clearance of virus infection is promoted by the interplay of neutralising antibodies with CD8^+^ T cells that eliminate intracellular virus reservoirs. CD8^+^ T cells may also reduce the influx of monocytes into infected lung tissue, which can be associated with undesirable IL-6 and TNF production and impaired antigen presentation^28,29^. The contributions of the immune effector systems to human protection from SARS-CoV-2 is not yet understood. Therefore, it appears prudent to develop COVID-19 vaccines that enlist concomitant cognate B cell, CD4^+^ T cell, and CD8^+^ T-cell responses.

The immunogenicity of BNT162b2 in rhesus macaques paralleled its immunogenicity in mice. Seven days after Dose 2 of 100 μg, the neutralising GMT reached 18-times that of a human SARS-CoV-2 convalescent serum panel and remained 3.3-times higher than this benchmark five weeks after the last immunisation. The strongly T_H_1-biased CD4^+^ T-cell response and IFNγ^+^ CD8^+^ T-cell response to BNT162b2 is a pattern favoured for vaccine safety and efficacy, providing added reassurance for clinical translation^30^. BNT162b2 protected 2-4 year old rhesus macaques from infectious SARS-CoV-2 challenge, with reduced detection of viral RNA in immunised animals compared to those that received saline and with no evidence of clinical exacerbation. Strong RT-qPCR evidence for lower respiratory tract protection was demonstrated by the absence of detectable SARS-CoV-2 RNA in serial BAL samples obtained starting 3 days after challenge of BNT162b2-immunised rhesus macaques.

We recently presented data from immunisation with BNT162b1, a vaccine candidate that has the same LNP-formulated m1Ψ nucleoside-modified RNA platform but expresses a trimerised, secreted RBD (Vogel et al., manuscript in preparation). The pattern, magnitude and durability of humoral and cellular responses to BNT162b1 in mice and macaques were in the range of those elicited by BNT162b2, as was protection of macaques from virus challenge, indicating that these features are largely class-intrinsic for this particular vaccine platform. BNT162b1 elicits high SARS-CoV-2 neutralizing titers and strong T_H_1-biased CD4^+^ and IFNγ^+^ and IL-2^+^ CD8^+^ T cell responses in humans, consistent with the preclinical findings^15,31,32^.

The selection of BNT162b2 over BNT162b1 for further clinical testing was largely driven by greater tolerability of BNT162b2 with comparable immunogenicity in clinical trials^15^ and the broader range of T-cell epitopes on the much larger full length spike. A global, pivotal, phase 3 safety and efficacy study of immunisation with BNT162b2 (NCT04368728) is now well under way and may answer those open questions that cannot be addressed by preclinical models.

## Materials and Methods

### Ethics statement

All mouse studies were performed at BioNTech SE, and protocols were approved by the local authorities (local welfare committee), conducted according to FELASA recommendations and in compliance with the German Animal Welfare Act and Directive 2010/63/EU. Only animals with an unobjectionable health status were selected for testing procedures.

Immunisations for the non-human primate (NHP) study were performed at the University of Louisiana at Lafayette-New Iberia Research Center (NIRC), which is accredited by the Association for Assessment and Accreditation of Laboratory Animal Care (AAALAC, Animal Assurance #: 000452). The work was in accordance with USDA Animal Welfare Act and Regulations and the NIH Guidelines for Research Involving Recombinant DNA Molecules, and Biosafety in Microbiological and Biomedical Laboratories. All procedures performed on these animals were in accordance with regulations and established guidelines and were reviewed and approved by an Institutional Animal Care and Use Committee or through an ethical review process. Infectious SARS-CoV-2 challenge for the NHP study was performed at the Southwest National Primate Research Center. Animal husbandry followed standards recommended by AAALAC International and the NIH Guide for the Care of Use of Laboratory Animals. This study was approved by the Texas Biomedical Research Institute Animal Care and Use Committee.

### Protein and peptide reagents

Purified recombinant SARS-CoV-2 S1 subunit including a histidine tag and the RBD tagged with the Fc region of human IgG1 (both Sino Biological) were used in ELISA to detect SARS-CoV-2 S-specific IgG in mice. Purified recombinant SARS-CoV-2 S1 and RBD with a histidine tag (both Sino Biological) were used for surface plasmon resonance (SPR) spectroscopy. An overlapping 15-mer peptide pool of the S protein was used for ELISpot, cytokine profiling and intracellular cytokine staining. An irrelevant peptide control (SPSYVYHQF, derived from gp70 AH-1^33^) or a CMV peptide pool was used as control for ELISpot assays. All peptides were obtained from JPT Peptide Technologies.

### Human convalescent sera

Human COVID-19 convalescent sera (n=38) were drawn from donors 18-83 years of age at least 14 days after PCR-confirmed diagnosis and at a time when the participants were asymptomatic. Serum donors had symptomatic infections (35/38), or had had been hospitalised (1/38). Sera were obtained from Sanguine Biosciences (Sherman Oaks, CA), the MT group (Van Nuys, CA) and Pfizer Occupational Health and Wellness (Pearl River, NY) and used across different studies as reference benchmark.

### Cell culture

Human embryonic kidney (HEK)293T/17 and Vero 76 cells (both ATCC) were cultured in Dulbecco’s modified Eagle’s medium (DMEM) with GlutaMAX™ (Gibco) supplemented with 10% fetal bovine serum (FBS [Sigma-Aldrich]). Cell lines were tested for mycoplasma contamination after receipt, before expansion and cryopreservation. For studies including NHP samples, Vero 76 and Vero CCL81 (both ATCC) cells were cultured in DMEM (Gibco) containing 2% HyClone fetal bovine and 100 U/mL penicillium/streptomycin (Gibco). Expi293F™ cells were grown in Expi293™ media and transiently transfected using ExpiFectamine™293 (all from Thermo Fisher Scientific).

### *In vitro* transcription and purification of RNA

To generate the template for RNA synthesis, a DNA fragment encoding the SARS-CoV-2 P2 S protein (based on GenBank: MN908947), including the amino acid exchanges K986P and V987P, was cloned into a starting plasmid vector with backbone sequence elements for improved RNA stability and translational efficiency^19,34^. Non-coding backbone elements included the regions from the T7 promoter to the 5’ and 3’ UTR plus a poly(A) tail (100 nucleotides) interrupted by a linker (A30LA70, 10 nucleotides). The DNA was purified, spectrophotometrically quantified, and *in vitro* transcribed by T7 RNA polymerase in the presence of a trinucleotide cap1 analogue ((m_2_^7,3’-O^)Gppp(m^2’-O^)ApG; TriLink) and of N^1^-methylpseudouridine-5’-triphosphate (m1ΨTP; Thermo Fisher Scientific) instead of uridine-5’-triphosphate (UTP)^35^. RNA was purified using magnetic particles^36^, integrity assessed by microfluidic capillary electrophoresis (Agilent Fragment Analyser), and concentration, pH, osmolality, endotoxin level and bioburden determined.

### Lipid-nanoparticle formulation of the RNA

Purified RNA was formulated into LNPs using an ethanolic lipid mixture of ionisable cationic lipid and transferred into an aqueous buffer system via diafiltration to yield an LNP composition similar to one previously described^37^. BNT162b2 was stored at −70 °C at a concentration of 0.5 mg/mL.

### mRNA transfection and P2 S translation

HEK293T/17 cells were transfected with 1 μg RiboJuice transfection reagent-mixed BNT162b2 RNA or with BNT162b2 (BNT162b2 RNA formulated as LNP) by incubation for 18 hours. Non-LNP formulated mRNA was diluted in Opti-MEM medium (Thermo Fisher Scientific) and mixed with the transfection reagents according to the manufacturer’s instructions (RiboJuice, Merck Millipore). Transfected HEK293T/17 cells were stained with Fixable Viability Dye (eBioscience). After fixation (Fixation Buffer, BioLegend), cells were permeabilised (Perm Buffer, eBioscience) and stained with a monoclonal SARS-CoV-2 spike S1 antibody (SinoBiological). Cells were acquired on a FACSCanto II flow cytometer (BD Biosciences) using BD FACSDiva software version 8.0.1 and analysed by FlowJo software version 10.6.2 (FlowJo LLC, BD Biosciences).

### P2 S expression and purification

To express P2 S for structural characterisation, a gene encoding the full length of SARS-CoV-2 (GenBank: MN908947) with two prolines substituted at residues 986 and 987 followed with a C-terminal HRV3C protease site and a TwinStrep tag was cloned into a modified pcDNA3.1(+) vector with the CAG promoter. The TwinStrep-tagged P2 S was expressed in Expi293 cells. Purification of the recombinant protein was based on a procedure described previously, with minor modifications^5^. Upon cell lysis, P2 S was solubilized in 1% NP-40 detergent. The TwinStrep-tagged protein was then captured with StrepTactin Sepharose HP resin in 0.5% NP-40. P2 S was further purified by size-exclusion chromatography and eluted as three distinct peaks in 0.02 % NP-40 as previously reported^5^. Peak 1, which consists of intact P2 S migrating at around 150 kDa, as well as dissociated S1 and S2 subunits, which co-migrate at just above 75 kDa, was used in the structural characterisation. Spontaneous dissociation of the S1 and S2 subunits mostly occurs throughout the course of the protein purification, starting at the point of detergent-mediated protein extraction.

### Biolayer interferometry

The binding of detergent NP-40 solubilized, purified P2 S to human ACE2 peptidase domain (ACE2 PD) and human neutralising monoclonal antibody B38^22^ was performed on Octet RED384 (FortéBio) at 25 °C in a running buffer (RB) consisting of 25 mM Tris pH7.5, 150 mM NaCl, 1 mM EDTA and 0.02% NP-40. Avi-tagged ACE2-PD was captured on streptavidin coated sensors and B38 antibody was captured on sensors coated with protein G. After initial baseline equilibration of 120 s, the sensors were dipped in 10 μg/mL solution of Avi-tagged ACE2-PD or B38 mAb for 300 s to achieve capture levels of 1 nM using the threshold function. The sensors were dipped in RB for 120 s for collecting baseline before they were dipped in a concentration series of purified P2 S samples for 300 s (association phase). The sensors were immersed in RB for measuring 600 s (dissociation phase). Data were reference subtracted and fit to a 1:1 binding model with R^2^ value greater than 0.95, to determine kinetics and affinity of binding, using Octet Data Analysis Software v10.0 (FortéBio).

### Cryo-electron microscopy sample preparation, data collection and data processing

For TwinStrep-tagged P2 S, 4 μL purified protein at 0.5 mg/mL were applied to gold Quantifoil R1.2/1.3 300 mesh grids freshly overlaid with graphene oxide. Sample was blotted using a Vitrobot Mark IV for 4 s with a force of −2 before being plunged into liquid ethane cooled by liquid nitrogen. 27,701 micrographs were collected from a two identically prepared grids on a Titan Krios operating at 300 keV equipped with a Gatan K2 Summit direct electron detector in super-resolution mode at a magnification of 165,000x, for a magnified pixel size of 0.435 Å at the specimen level. Data were collected from each grid over a defocus range of −1.2 to −3.4 μm with a total electron dose of 50.32 and 50.12 e^−^/Å^2^, respectively, fractionated into 40 frames over a 6-second exposure for 1.26 and 1.25 e^−^/Å^2^/frame. On-the-fly motion correction, CTF estimation, and particle picking and extraction with a box size of 450 pixels were performed in Warp^38^, during which super-resolution data were binned to give a pixel size of 0.87 Å. A total of 1,119,906 particles were extracted. All subsequent processing was performed in RELION 3.1-beta^39^. Particle heterogeneity was filtered out with 2D and 3D classification to filter out bad particles, yielding a set of 73,393 particles, which refined to 3.6 Å with C3 symmetry. 3D classification of this dataset without particle alignment separated out one class with a single RBD up, representing 15,098 particles. The remaining 58,295 particles, in three RBD ‘down’ conformation, were refined to give a final model at 3.29 Å. The atomic model from PDB ID 6XR8^5^ was rigid-body fitted into the map density, then flexibly fitted to the density using real-space refinement in Phenix^40^ alternating with manual building in Coot^41^. The cryo-EM model validation is provided in Extended Data Fig 3b, the full cryo-EM data processing workflow in Extended Data Fig. 3c, and the model refinement statistics in Extended Data Table 1.

### Immunisation

#### Mice

Female BALB/c mice (Janvier; 8-12 weeks) and were randomly allocated to groups. BNT162b2 was diluted in PBS, 300 mM sucrose or saline (0.9% NaCl) and injected IM into the gastrocnemius muscle at a volume of 20 μL under isoflurane anaesthesia.

#### Rhesus macaques (Macaca mulatta)

Male rhesus macaques (2–4 years) were randomly assigned to receive either BNT162b2 or saline placebo control in 0.5 mL volume administered by IM injection in the left quadriceps muscle on Days 0 and 21.

### Tissue preparation

#### Mice

Peripheral blood was collected from the retro-orbital venous plexus under isoflurane anaesthesia or *vena facialis* without prior anesthetisation. Blood was centrifuged for 5 minutes at 16.000 × g, and the serum was immediately used for downstream assays or stored at −20 °C. Spleen single-cell suspensions were prepared in PBS by mashing tissue against the surface of a 70 μm cell strainer (BD Falcon). Erythrocytes were removed by hypotonic lysis. Popliteal, inguinal and iliac lymph nodes were pooled, cut into pieces, digested with collagenase D (1 mg/mL; Roche) and passed through cell strainers.

#### Rhesus macaques (Macaca mulatta)

Serum was obtained before immunisation and on Days 14, 21, 28, 35, 42, and 56. PBMCs were obtained before immunisation and on Days 7, 28, and 42, except that PBMCs were not obtained from the buffer-immunised group on Day 28. Blood for serum and PBMCs was collected in compliance with animal protocol 2017-8725-023 approved by the NIRC Institutional Animal Care and Use Committee. Animals were anesthetised with ketamine HCl (10 mg/kg; IM) during blood collection and immunisation, and monitored for adequate sedation.

### S1- and RBD-binding IgG assay

For mouse sera, MaxiSorp plates (Thermo Fisher Scientific) were coated with recombinant S1 or RBD (100 ng/100 μL) in sodium carbonate buffer, and bound IgG was detected using an HRP-conjugated secondary antibody and TMB substrate (Biotrend). Data collection was performed using a BioTek Epoch reader and Gen5 software version 3.0.9. For concentration analysis, the signal of the specific samples was correlated to a standard curve of an isotype control. For rhesus macaque and human sera, a recombinant SARS-CoV-2 S1 containing a C-terminal Avitag™ (Acro Biosystems) was bound to streptavidin-coated Luminex microspheres. Bound rhesus macaque or human anti-S1 antibodies present in the serum were detected with a fluorescently labelled goat anti-human polyclonal secondary antibody (Jackson ImmunoResearch). Data were captured as median fluorescent intensities (MFIs) using a Bioplex200 system (Bio-Rad) and converted to U/mL antibody concentrations using a reference standard consisting of 5 pooled human COVID-19 convalescent serum samples (obtained >14 days PCR diagnosis), diluted in antibody depleted human serum with arbitrary assigned concentrations of 100 U/mL and accounting for the serum dilution factor.

### Binding kinetics of antigen-specific IgGs using surface plasmon resonance spectroscopy

Binding kinetics of murine S1- and RBD-specific serum IgGs was determined using a Biacore T200 device (Cytiva) with HBS-EP running buffer (BR100669, Cytiva) at 25 °C. Carboxyl groups on the CM5 sensor chip matrix were activated with a mixture of 1-ethyl-3-(3-dimethylaminopropyl) carbodiimidehydrochloride (EDC) and N-hydroxysuccinimide (NHS) to form active esters for the reaction with amine groups. Anti-mouse-Fc-antibody (Jackson ImmunoResearch) was diluted in 10 mM sodium acetate buffer pH 5 (30 μg/mL) for covalent coupling to immobilisation level of ~10,000 response units (RU). Free N-hydroxysuccinimide esters on the sensor surface were deactivated with ethanolamine.

Mouse serum was diluted 1:50 in HBS-EP buffer and applied at 10 μL/min for 30 seconds to the active flow cell for capture by immobilised antibody, while the reference flow cell was treated with buffer. Binding analysis of captured murine IgG antibodies to S1-His or RBD-His (Sino Biological) was performed using a multi-cycle kinetic method with concentrations ranging from 25 to 400 nM or 1.5625 to 50 nM, respectively. An association period of 180 seconds was followed by a dissociation period of 600 seconds with a constant flow rate of 40 μL/min and a final regeneration step. Binding kinetics were calculated using a global kinetic fit model (1:1 Langmuir, Biacore T200 Evaluation Software Version 3.1, Cytiva).

### VSV-SARS-CoV-2 spike variant pseudovirus neutralisation

A recombinant replication-deficient vesicular stomatitis virus (VSV) vector that encodes GFP instead of VSV-G (VSVΔG-GFP) was pseudotyped with SARS-CoV-2 S protein according to published pseudotyping protocols^42,43^. In brief, HEK293T/17 monolayers transfected to express SARS-CoV-2 S truncated of the C-terminal cytoplasmic 19 amino acids (SARS-CoV-2-S-CΔ19) were inoculated with VSVΔG-GFP vector. After incubation for 1 hour at 37 °C, the inoculum was removed and cells were washed with PBS before medium supplemented with anti-VSV-G antibody (clone 8G5F11, Kerafast Inc.) was added to neutralise residual input virus. VSV/SARS-CoV-2 pseudovirus-containing medium was harvested 20 hours after inoculation, 0.2 μm filtered and stored at −80 °C.

Serial dilutions of mouse serum samples were prepared and pre-incubated for 10 minutes at room temperature with VSV/SARS-CoV-2 pseudovirus suspension (4.8 × 10^3^ infectious units [IU]/mL) before transferring the mix to Vero 76 cells. Inoculated Vero-76 cells were incubated for 20 hours at 37 °C. Plates were placed in an IncuCyte Live Cell Analysis system (Sartorius) and incubated for 30 minutes prior to the analysis (IncuCyte 2019B Rev2 software). Whole well scanning for brightfield and GFP fluorescence was performed using a 4× objective. The 50% pseudovirus neutralisation titre (pVNT_50_) was reported as the reciprocal of the first serum dilution yielding a 50% reduction in GFP-positive infected cell number per well compared to the mean of the no serum pseudovirus positive control. Each serum sample dilution was tested in duplicates.

### IFNγ and IL-4 ELISpot

Murine ELISpot assays were performed with mouse IFNγ ELISpot^PLUS^ kits according to the manufacturer’s instructions (Mabtech). A total of 5 × 10^5^ splenocytes was *ex vivo* restimulated with the full-length S peptide mix (0.1 μg/mL final concentration per peptide) or controls (gp70-AH1 [SPSYVYHQF]^33^, 4 μg/mL; Concanavalin A [ConA], 2 μg/mL [Sigma]). Streptavidin-alkaline phosphatase (ALP) and BCIP/NBT-plus substrate were added, and spots counted using an ELISpot plate reader (ImmunoSpot^®^ S6 Core Analyzer [CTL]). Spot numbers were evaluated using ImmunoCapture Image Aquision Software V7.0 and ImmunoSpot 7.0.17.0 Professional. Spot counts denoted too numerous to count by the software were set to 1,500. For T-cell subtyping, CD8^+^ T cells and CD4^+^ T cells were isolated from splenocyte suspensions using MACS MicroBeads (CD8a [Ly-2] and CD4 [L3T4] [Miltenyi Biotec]) according to the manufacturer’s instructions. 1 × 10^5^ CD8^+^ or CD4^+^ T cells were subsequently restimulated with 5 × 10^4^ syngeneic bone marrow-derived dendritic cells loaded with full-length S peptide mix (0.1 μg/mL final concentration per peptide) or cell culture medium as control. Purity of isolated T-cell subsets was determined by flow cytometry.

Rhesus macaque PBMCs were tested with commercially available NHP IFNγ and IL-4 ELISpot assay kits (Mabtech, Sweden). Cryopreserved rhesus macaque PBMCs were thawed in pre-warmed AIM-V media (Thermo Fisher Scientific, US) with Benzonase (EMD Millipore, US). For IFNγ ELISpot, 1.0 × 10^5^ PBMCs and 2.5 × 10^5^ PBMCs for IL-4 ELISpot were stimulated *ex vivo* with 1 μg/mL of the full-length S overlapping peptide mix. Tests were performed in triplicate wells and media-DMSO, a CMV peptide pool and PHA (Sigma) were included as controls. After 24 hours for IFNγ and 48 hours for IL-4, Streptavidin-HRP and AEC substrate (BD Bioscience) were added, and spots counted using a CTL ImmunoSpot S6 Universal Analyzer (CTL, US). Results shown are background (Media-DMSO) subtracted and normalized to SFC/10^6^ PBMCs.

### Flow cytometry for analysis of cell mediated immunity

For mouse T-cell analysis in peripheral blood, erythrocytes from 50 μL freshly drawn blood were lysed (ACK lysing buffer [Gibco]), and cells were stained with Fixable Viability Dye (eBioscience) and primary antibodies in the presence of Fc block in flow buffer (DPBS [Gibco] supplemented with 2% FCS, 2 mM EDTA [both Sigma] and 0.01% sodium azide [Morphisto]). After staining with secondary biotin-coupled antibodies in flow buffer, cells were stained extracellularly against surface markers with directly labelled antibodies and streptavidin in Brilliant Stain Buffer Plus (BD Bioscience) diluted in flow buffer. Cells were washed with 2% RotiHistofix (Carl Roth), fixed (Fix/Perm Buffer, FoxP3/Transcription Factor Staining Buffer Set [eBioscience]) and permeabilised (Perm Buffer, FoxP3/Transcription Factor Staining Buffer Set [eBioscience]) overnight. Permeabilised cells were intracellularly treated with Fc block and stained with antibodies against transcription factors in Perm Buffer.

For mouse T-cell analysis in lymphoid tissues, 1.5 × 10^6^ lymph node and 4 × 10^6^ spleen cells were stained for viability and extracellular antigens with directly labelled antibodies. Fixation, permeabilisation and intracellular staining was performed as described for blood T-cell staining. For mouse B-cell subtyping in lymphoid tissues, 2.5 × 10^5^ lymph node and 1 × 10^6^ spleen cells were treated with Fc block, stained for viability and extracellular antigens as described for blood T-cell staining and fixed with 2% RotiHistofix overnight.

For mouse intracellular cytokine staining in T cells, 4 × 10^6^ spleen cells were *ex vivo* restimulated with 0.5 μg/mL final concentration per peptide of full-length S peptide mix or cell culture medium (no peptide) as control in the presence of GolgiStop and GolgiPlug (both BD Bioscience) for 5 hours. Cells were stained for viability and extracellular antigens as described for lymphoid T-cell staining. Cells were fixed with 2% RotiHistofix and permeabilised overnight. Intracellular staining was performed as described for blood T-cell staining.

Mouse cells were acquired on a BD Symphony A3 or BD Celesta (B-cell subtyping) flow cytometer (BD Bioscience) using BD FACSDiva software version 9.1 or 8.0.1.1, respectively, and analysed with FlowJo 10.6 (FlowJo LLC, BD Biosciences).

For rhesus macaques intracellular cytokine staining in T cells, 1.5 × 10^6^ PBMCs were stimulated with the full-length S peptide mix at 1 μg/mL, Staphyloccocus enterotoxin B (SEB; 2 μg/mL) as positive control, or 0.2% DMSO as negative control. GolgiStop and GolgiPlug (both BD Bioscience) were added. Following 37 °C incubation for 12 to 16 h, cells were stained for viability and extracellular antigens after blocking Fc binding sites with directly labelled antibodies. Cells were then fixed and permeabilized with BDCytoFix/CytoPerm solution (BD Bioscience), intracellular staining was performed in perm buffer for 30 min at RT. Cells were washed, resuspended in 2% FBS/PBS buffer and acquired on a LSR Fortessa. Data analyzed by FlowJo (10.4.1). Results shown are background (Media-DMSO) subtracted.

### Cytokine profiling

Mouse splenocytes were re-stimulated for 48 hours with full-length S peptide mix (0.1 μg/mL final concentration per peptide) or cell culture medium (no peptide) as control. Concentrations of IFNγ, IL-2, IL-4, IL-5 and IL-13 in supernatants were determined using a bead-based, 11-plex T_H_1/T_H_2 mouse ProcartaPlex multiplex immunoassay (Thermo Fisher Scientific) according to the manufacturer’s instructions. Fluorescence was measured with a Bioplex200 system (Bio-Rad) and analysed with ProcartaPlex Analyst 1.0 software (Thermo Fisher Scientific). Values below the lower limit of quantification (LLOQ) were set to zero.

### SARS-CoV-2 neutralisation

The SARS-CoV-2 neutralisation assay used a previously described strain of SARS-CoV-2 (USA_WA1/2020) that had been rescued by reverse genetics and engineered by the insertion of an mNeonGreen (mNG) gene into open reading frame 7 of the viral genome^25^. This reporter virus generates similar plaque morphologies and indistinguishable growth curves from wild-type virus. Viral master stocks were grown in Vero 76 cells as previously described^44^. When testing human convalescent serum specimens, the fluorescent neutralisation assay produced comparable results as the conventional plaque reduction neutralisation assay. Serial dilutions of heat-inactivated sera were incubated with the reporter virus (2 × 10^4^ PFU per well) to yield approximately a 10-30% infection rate of the Vero CCL81 monolayer for 1 hour at 37 °C before inoculating Vero CCL81 cell monolayers (targeted to have 8,000 to 15,000 cells in the central field of each well at the time of seeding, one day before infection) in 96-well plates to allow accurate quantification of infected cells. Cell counts were enumerated by nuclear stain (Hoechst 33342) and fluorescent virally infected foci were detected 16-24 hours after inoculation with a Cytation 7 Cell Imaging Multi-Mode Reader (Biotek) with Gen5 Image Prime version 3.09. Titers were calculated in GraphPad Prism version 8.4.2 by generating a 4-parameter (4PL) logistical fit of the percent neutralisation at each serial serum dilution. The 50% neutralisation titre (VNT_50_) was reported as the interpolated reciprocal of the dilution yielding a 50% reduction in fluorescent viral foci.

### SARS-CoV-2 challenge of rhesus macaques (*Macaca mulatta*)

The SARS-CoV-2 inoculum was obtained from a stock of 2.1 × 10^6^ PFU/mL previously prepared at Texas Biomedical Research Institute (San Antonio, TX), aliquoted into single use vials, and stored at −70 °C. The working virus stock was generated from two passages of the SARS-CoV-2 USA-WA1/2020 isolate (a 4^th^ passage seed stock purchased from BEI Resources; NR-52281) in Vero 76 cells. The virus was confirmed to be SARS-CoV-2 by deep sequencing and identical to the published sequence (GenBank accession number MN985325.1).

BNT162b2-immunised (n=6) and age-matched saline control-immunised (n=3) male rhesus macaques (control) were challenged with 1.05 × 10^6^ plaque forming units of SARS-CoV-2 USA-WA1/2020 isolate, split equally between the intranasal (IN; 0.25 mL) and intratracheal (IT; 0.25 mL) routes as previously described^26^. The challenge was performed 55 days after the second immunisation. A separate sentinel group of non-immunised age- and sex-matched animals (n=3) was mock challenged with DMEM supplemented with 10% FCS IN (0.25 mL) and IT (0.25 mL). Approximately two weeks prior to challenge, animals were moved to the ABSL-3 facility at Southwest National Primate Research Center (SNPRC; San Antonio, TX). Animals were monitored regularly by a board-certified veterinary clinician for rectal body temperature, weight and physical examination. Specimen collection was performed under tiletamine zolazepam (Telazol) anaesthesia as described^26^. Nasal and oropharyngeal swabs were collected from all macaques at Day 0, 1, 3, and 6 (relative to the day of challenge), from BNT162b2-immunised macaques on Day 7 or 8, and from control and sentinel macaques on Day 10. Bronchoalveolar lavage (BAL) was performed on all macaques the week before challenge and on Days 3 and 6 post-challenge and on BNT162b2-immunised macaques on Day 7 or 8. BAL was performed by instilling four times 20 mL of saline. These washings were pooled, aliquoted and stored frozen at −70 °C. Necropsy was performed on BNT162b2-immunised animals on Day 7 or 8. Control and sentinel animals were not necropsied to allow re-challenge (control) or challenge (sentinel) on a subsequent day.

### Reverse-transcription quantitative polymerase chain reaction

To detect and quantify SARS-CoV-2 in NHP, viral RNA was extracted from nasal swabs, OP swabs, and BAL specimens as previously described^45–47^ and tested by RT-qPCR as previously described^26^. Briefly, 10 μg yeast tRNA and 1 × 10^3^ PFU of MS2 phage (*Escherichia coli* bacteriophage MS2, ATCC) were added to each thawed sample, and RNA extraction performed using the NucleoMag Pathogen kit (Macherey-Nagel). The SARS-CoV-2 RT-qPCR was performed on extracted RNA using a CDC-developed 2019-nCoV_N1 assay on a QuantStudio 3 instrument (Applied Biosystems). The cut-off for positivity (limit of detection, LOD) was established at 10 gene equivalents (GE) per reaction (800 GE/mL). Samples were tested in duplicate. On day 6, one BAL specimen from the control group and one day 1 nasal swab from the BNT162b1-immunised group had, on repeated measurements, viral RNA levels on either side of the LLOD. These specimens were categorised as indeterminate and excluded from the graphs and the analysis.

### Statistics and reproducibility

No statistical methods were used to predetermine group and samples sizes (n). All experiments were performed once. P-values reported for RT-qPCR analysis were determined by nonparametric analysis (Friedman’s test) based on the ranking of viral RNA shedding data within each day. PROC RANK and PROC GLM from SAS^®^ 9.4 were used to calculate the p-values. Samples from post challenge days (Days 3, 6 and end of protocol [EOP] for BAL; Days 1, 3, 6 and 10 [control and sentinel] or EOP [BNT162b2-immunised] for nasal and oropharyngeal swabs) were included in the analysis. Indeterminate results were excluded from this analysis. All remaining analyses were carried out using GraphPad Prism 8.4.

### Data availability

The data that support the findings of this study are available from the corresponding author upon reasonable request.

## Acknowledgements

We thank T. Garretson for advice on NHP serology studies, R. F. Sommese and K. F. Fennell for technical assistance for molecular cloning and cell based binding. Valuable support and assistance from R. de la Caridad Güimil Garcia1, A. Su, M. Dvorak, M. Drude, F. Zehner, T. Lapin, B. Ludloff, S. Hinz, F. Bayer, J. Scholz, A.L. Ernst, T. Sticker and S. Wittig resulted in a rapid availability of oligonucleotides and DNA templates. E. Boehm, K. Goebel, R. Frieling, C. Berger, S. Koch, T. Wachtel, J. Leilich, M. Mechler, R. Wysocki, M. Le Gall, A. Czech and S. Klenk carried out RNA production and analysis. Without their commitment during this pandemic situation, this vaccine candidate could not have been transferred to non-clinical studies in light speed. B. Weber, J. Vogt, S. Krapp, K. Zwadlo, J. Mottl, J. Mühl and P. Windecker supported the mouse studies and serological analysis with excellent technical assistance. We thank A. Ota-Setlik for ELISpot testing of NHP samples. Radiologists A. K. Voges, M. R. Gutman and E. Clemmons advised on the radiographic scoring. We thank Polymun Scientific for excellent formulation services as well as Acuitas Therapeutics for fruitful discussions.

## Author Contributions

U.S. conceived and conceptualised the work and strategy. S.He., S.C.D., C.K. and M.C.G. designed primers and cloned all constructs. T.Z., S.F., J.S. and A.N.K. developed, planned, performed and supervised RNA synthesis and analysis. E.H.M. purified P2 S. P.V.S. developed and performed biolayer interferometry. J.A.L. and S.H. performed electron microscopy and solved the structure of the complex. Y.C. supervised the structural and biophysical characterisation and analysed the structures. A.M. and B.G.L. performed surface plasmon resonance spectroscopy. A.G. and S.A.K. planned, performed and analysed *in vitro* studies. F.B., T.K., C.R. managed formulation strategy. A.B.V., M.V., L.M.K. designed mouse studies and analysed and interpreted data. A.P., S.E., D.P. and G.S. performed and analysed the S1-binding IgG assays. R.C., Jr. and K.J.A. performed and analysed viral RT-qPCR data. A.M., B.S., A.W., C.F.-G. and P.-Y.S. performed and analysed pVNT and VNT assays. D.E., D.S., B. J., Y.F, H.J. performed *in vivo* studies and ELISpot assays. A.B.V., K.C.W. J.L., M.S.M. and M.V. planned, analysed and interpreted ELISpot assays. L.M.K., J.L., D.E., Y.F., H.J., A.P.H. M.S.M. and P.A planned, performed and analysed flow cytometry assays. A.B.V., L.M.K., Y.F. and H.J. planned, performed, analysed and interpreted cytokine release assays. I.K., K.A.S., K.T., D.K. and P.R.D. designed NHP studies and analysed and interpreted data. K.T., M.P., I.L.S. and W.K. oversaw NHP immunogenicity and serology testing. S.H.-U. and K.B. provided veterinary services for NHPs. J.C., T.C. and J.O managed the NHP colony. A.B.V., I.K., Y.C., A.M., M.V, L.M.K., C.T., K.S., Ö.T., P.R.D, K.U.J. and U.S. wrote the manuscript. All authors supported the review of the manuscript.

## Additional Information

Supplementary Information is available for this study.

Correspondence and requests for materials should be addressed to Ugur Sahin.

---

**Extended Data Figure 1.**
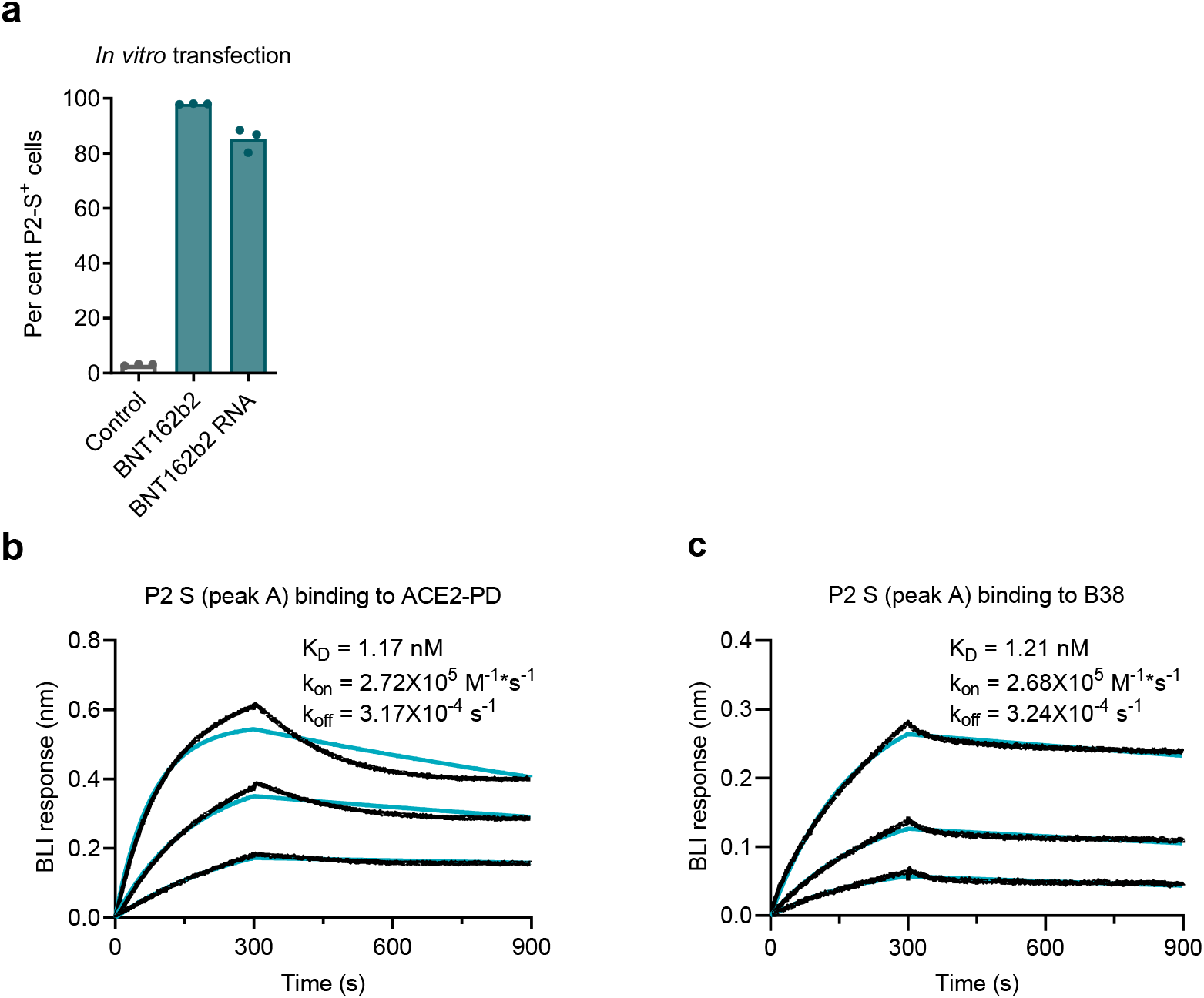
P2 S expression and receptor affinity. **a,** Fraction of HEK293T/17 cells expressing P2 S determined by flow cytometry after incubation with BNT162b2 RNA formulated as LNP (BNT162b2), BNT162b2 RNA formulated with a transfection reagent (BNT162b2 RNA), or no RNA (Control), determined by flow cytometry. **b, c,** P2 S with a C-terminal TwinStrep tag, expressed in Expi293F cells, was detergent solubilized and purified by affinity and size exclusion chromatography. Protein from the first peak of a size exclusion column, containing intact P2 S and dissociated S1 and S2 fragments, was assayed by biolayer interferometry. Sensorgram of the binding kinetics of TwinStrep-tagged P2 S to immobilised b, human ACE2-PD and c, B38 monoclonal antibody.

**Extended Data Figure 2.**
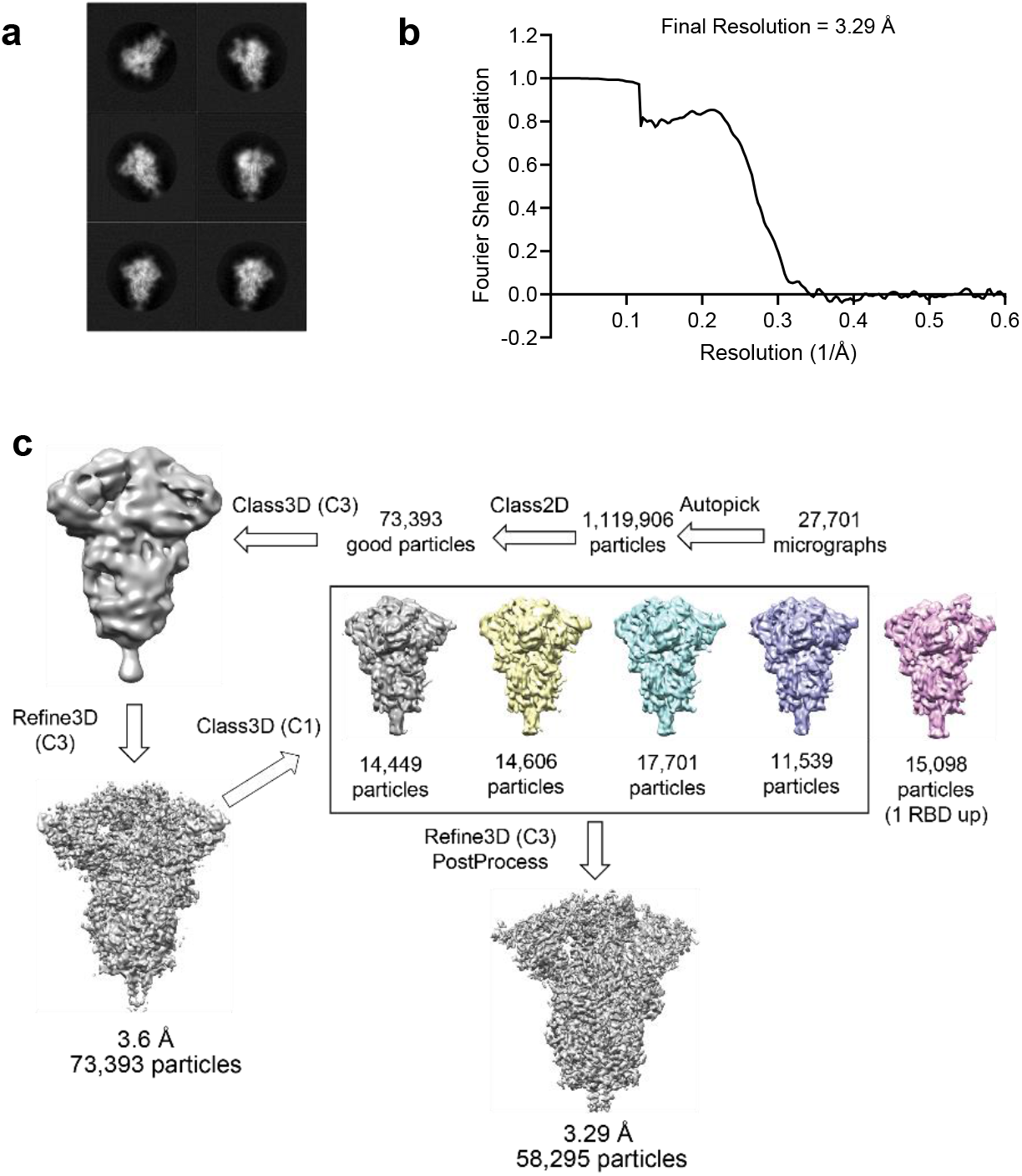
Structure analysis of BNT162b2-encoded P2 S by cryo-electron microscopy. **a**, 2D class averages of TwinStrep-tagged P2 S particles extracted from cryo-EM micrographs. Box edge: 39.2 nm in each dimension. **b**, Fourier shell correlation curve from RELION gold-standard refinement of the P2 S trimer. **c**, Flowchart for cryo-EM data processing of the complex.

**Extended Data Figure 3.**
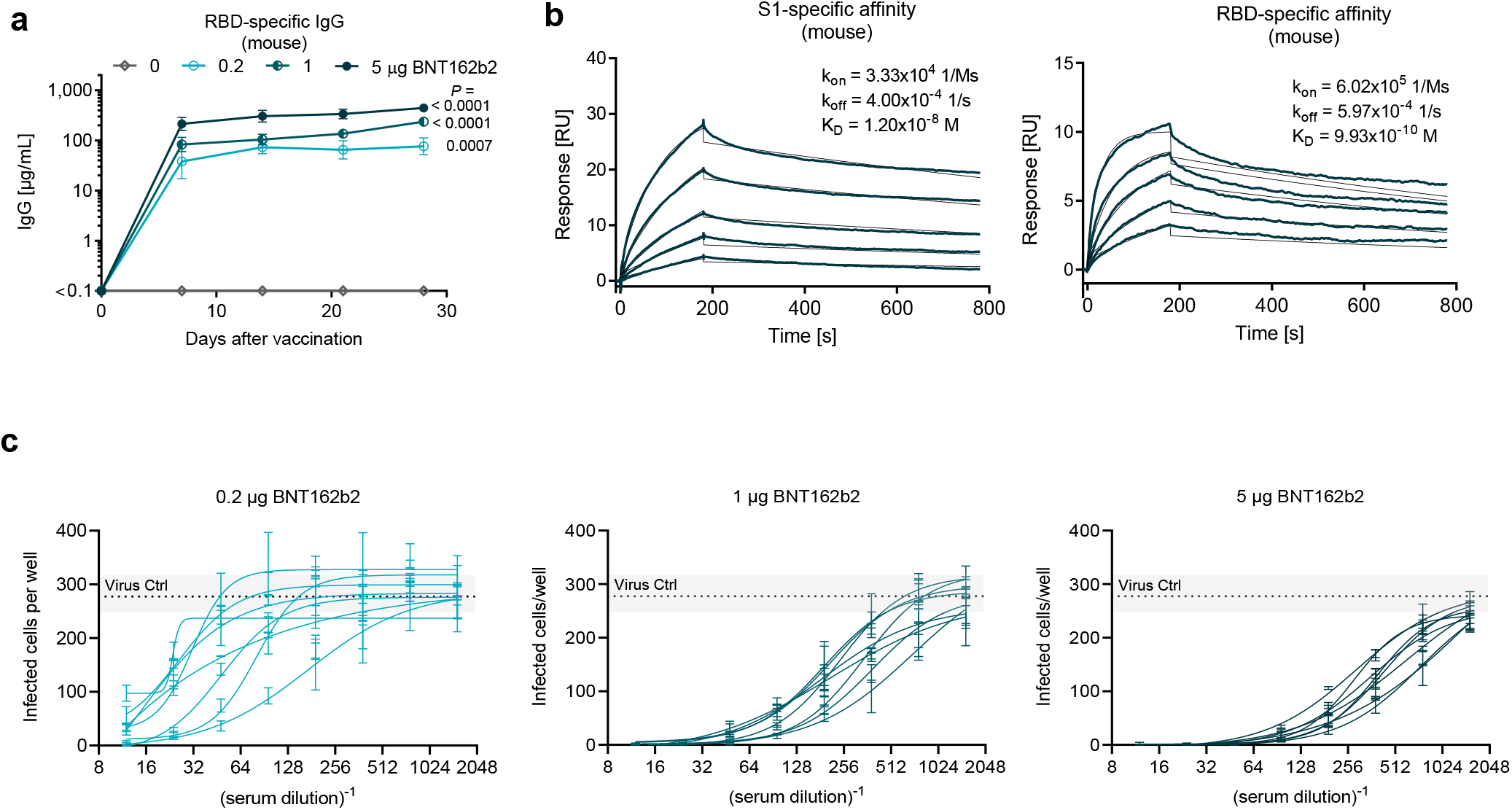
BNT162b2-elicited antibody responses in mice. BALB/c mice (n=8 per group) were immunised intramuscularly (IM) with 0.2, 1 or 5 μg of BNT162b2 or buffer. **a**, RBD-binding IgG responses in sera obtained 7, 14, 21 and 28 days after immunisation, determined by ELISA. For day 0, a pre-screening of randomised animals was performed (n=4). Geometric mean of each group is shown. **b**, Representative surface plasmon resonance sensorgram of the binding kinetics of recombinant S1 (*left*) and RBD (*right*) to immobilised mouse IgG from serum 28 days after immunisation with 5 μg BNT162b2 (n=8). Actual binding (dark blue) and the best fit of the data to a 1:1 binding model (thin line in black). **c**, Number of infected cells per well in pseudovirus neutralisation assays conducted with serially diluted mouse serum samples obtained 28 days after immunisation with BNT162b2 are shown (n=8 per group, see also Figure 2b).

**Extended Data Figure 4.**
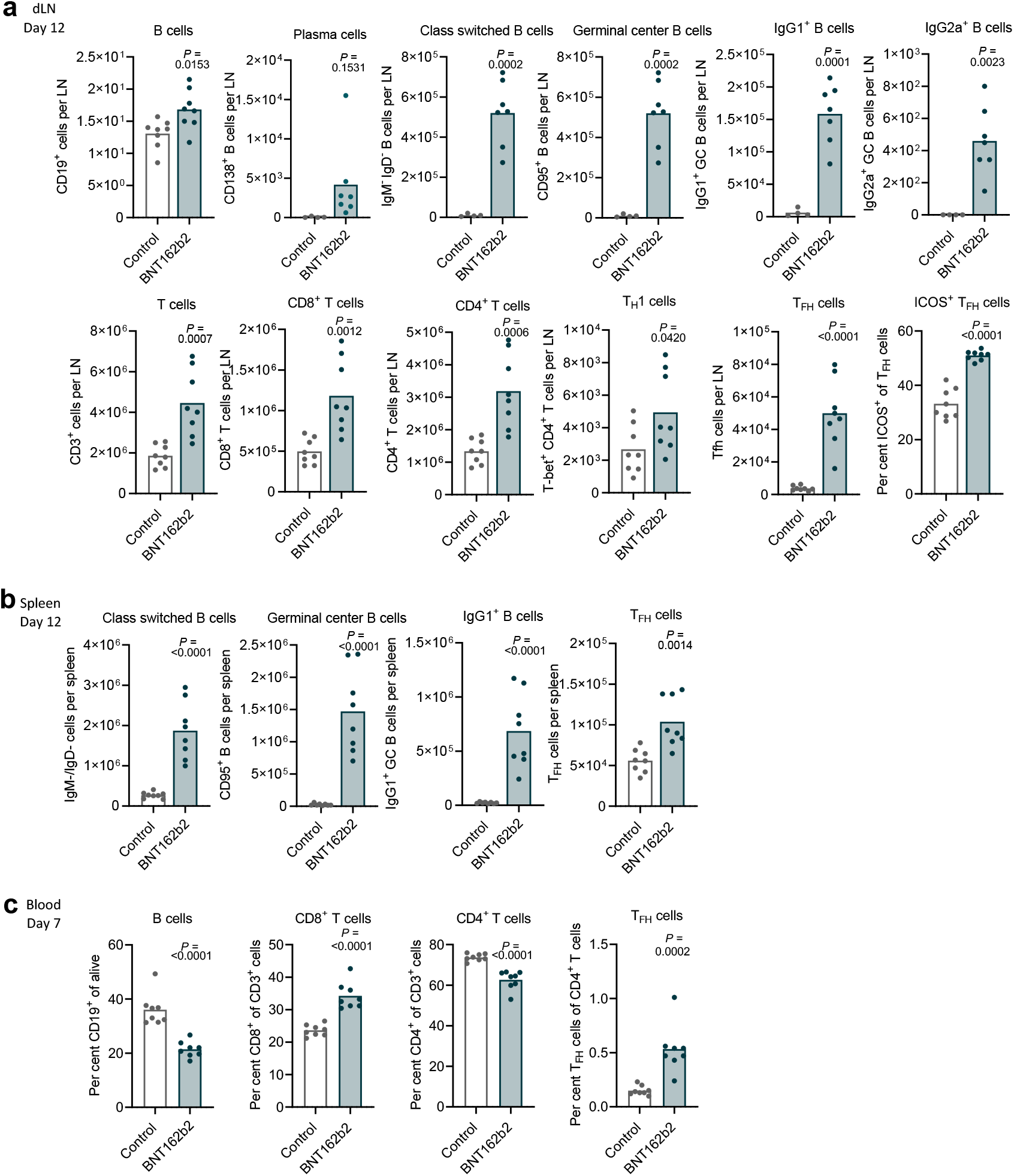
B-cell and T-cell phenotyping in lymph nodes, spleen and blood of BNT162b2 immunised mice. Mice (n=8 per group) were immunised with 5 μg BNT162b2 or buffer (Control). P-values were determined by an unpaired two-tailed t-test. **a**+**b**, B-cell and T-cell numbers 12 days after immunisation in (**a**) draining lymph nodes (dLN; for B-cell subtyping only: n=4 for control, n=7 for BNT162b2 immunised group) or (**b**) the spleen, determined by flow cytometry. The percentage of ICOS^+^ cells among T follicular helper cells (T_FH_) in dLNs is depicted on the lower right in (a). **c**, B- and T-cell fractions in the blood seven days after immunisation.

**Extended Data Figure 5.**
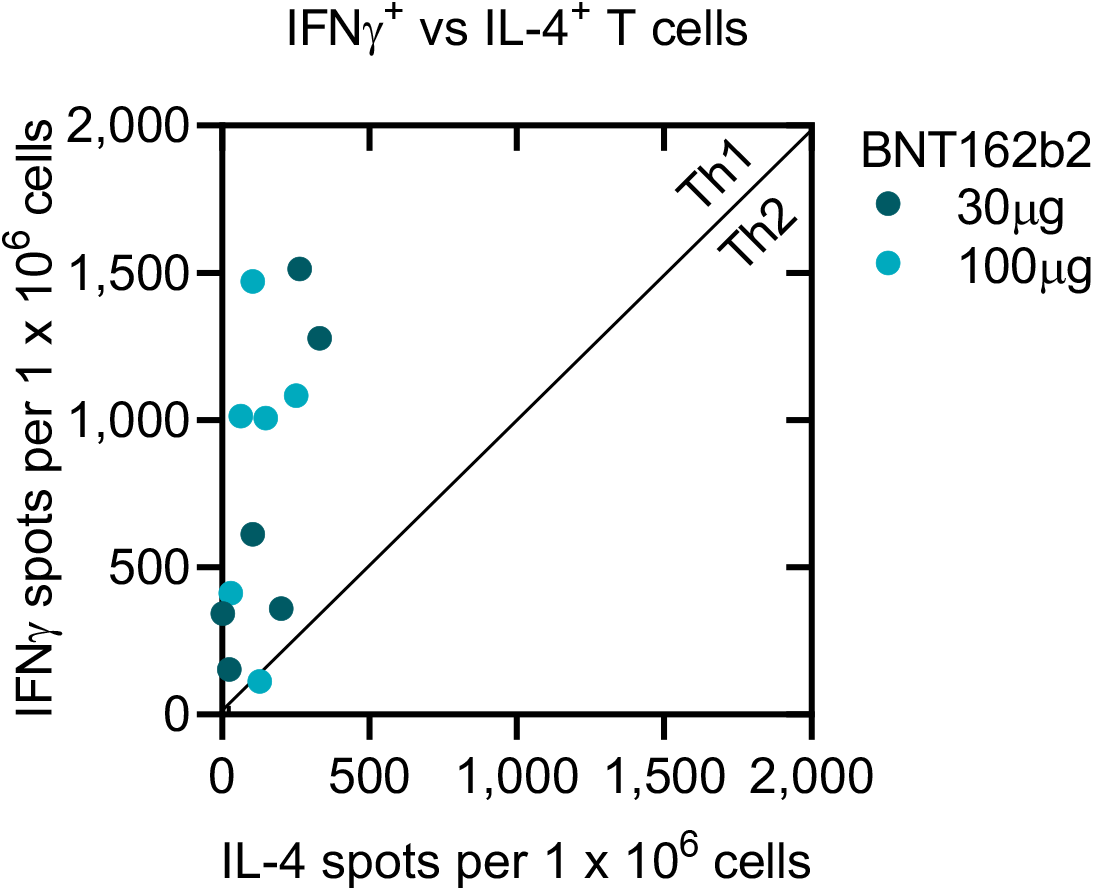
Scatterplot of IL-4 vs. IFNγ ELISpot of S peptide stimulated PBMCs collected on Day 42. Rhesus macaques (n=6 per group) were immunised on Days 0 and 21 with 30 μg or 100 μg BNT162b2 (see Fig. 3c,d) and individual animal values are shown.

**Extended Data Table 1.**
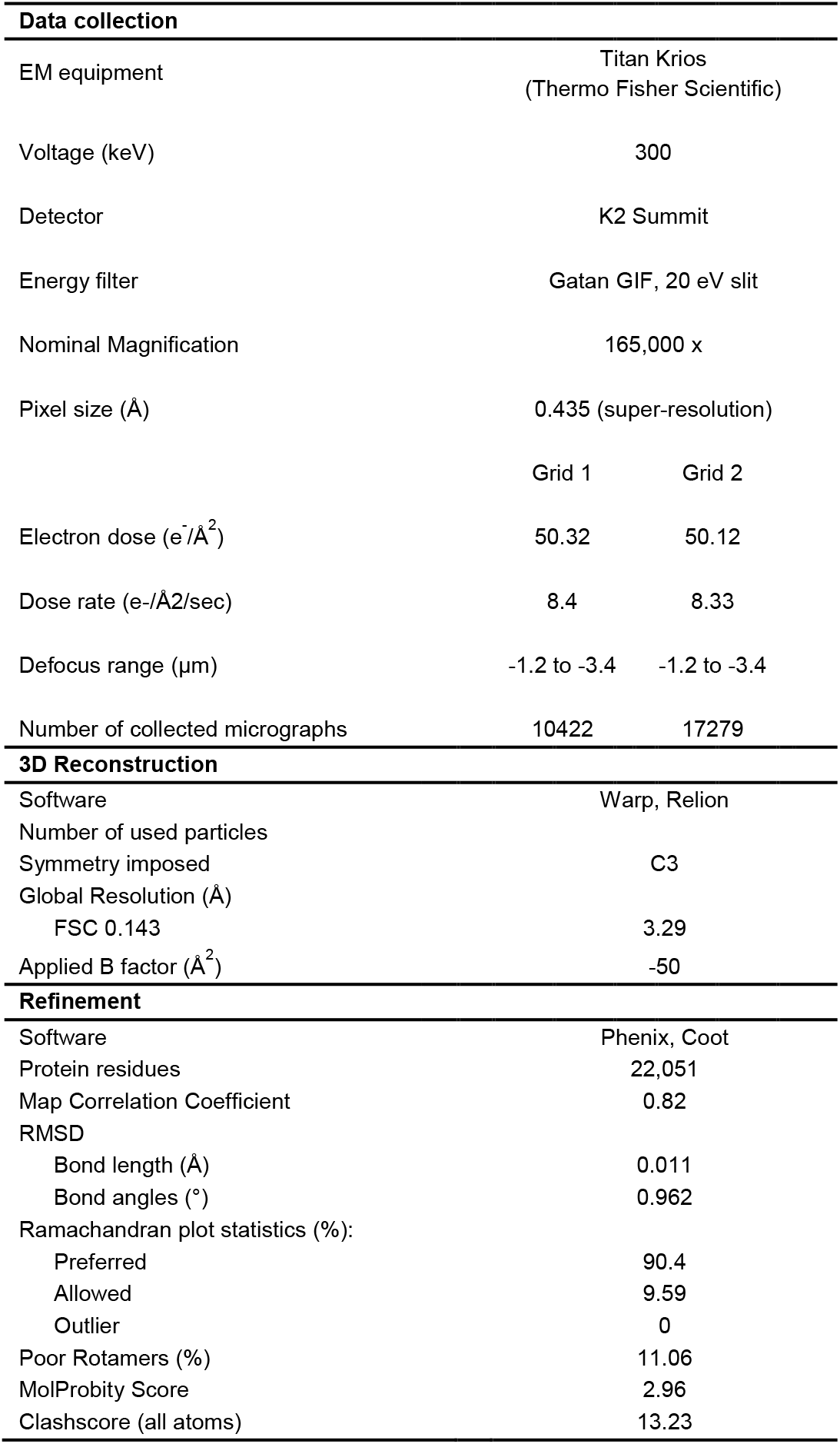
Electron cryomicroscopy data collection, 3D reconstruction and refinement statistics.

